# Microgravity modulates effects of chemotherapeutic drugs on cancer cell migration

**DOI:** 10.1101/2019.12.29.890632

**Authors:** Devika Prasanth, Sindhuja Suresh, Sruti Prathivadhi-Bhayankaram, Michael Mimlitz, Noah Zetocha, Bong Lee, Andrew Ekpenyong

## Abstract

Microgravity or the condition of apparent weightlessness causes bone, muscular and immune system dysfunctions in astronauts following spaceflights. These organ and system-level dysfunctions correlate with changes induced at the single cell level both by simulated microgravity on earth as well as microgravity conditions in outer space (as in the international space station). Reported changes in single bone cells, muscle cells and white blood cells include structural/morphological abnormalities, changes in gene expression, protein expression, metabolic pathways and signaling pathways, suggesting that cells mount some response or adjustment to microgravity. However, the implications of such adjustments on many cellular functions and responses are not clear largely because the primary mechanism of gravity sensing in animal cells is unknown. Here we used a rotary cell culture system developed by NASA, to subject leukemic and erythroleukemic cancer cells to microgravity for 48 hours and then quantified their innate immune-response to common anti-cancer drugs using biophysical parameters and our recently developed quantum-dots-based fluorescence spectroscopy. We found that leukemic cancer cells treated with daunorubicin show increased chemotactic migration (p < 0.01) following simulated microgravity (μg) compared to normal gravity on earth (1g). However, cells treated with doxorubicin showed enhanced migration both in 1g and following μg. Our results show that microgravity modulates cancer cell response to chemotherapy in a drug-dependent manner. These results suggest using simulated microgravity as an immunomodulatory tool for the development of new immunotherapies for both space and terrestrial medicine.

## 1. Introduction

Physical forces including electromagnetism and gravity have shaped the evolution of life on earth and continue to influence living processes in organisms. Changes in such forces produce profound biological effects. Microgravity in outer space and simulated microgravity on earth cause changes in biological cells, tissues and organs. In order to enhance space exploration, NASA and other space programs have developed several spaceflight analogue systems on earth, including unique suspension cell culture systems such as the rotary cell culture system, RCCS (1). It turns out that such analogue systems lead to discoveries and inventions with potential use for enhancing terrestrial life, medicine and further research. In particular, cancer research has employed microgravity as a condition for studying mechanisms that control cancer cell growth and function (2, 3). Recently reported effects of simulated microgravity on cancer cells include increase in migration of non-small cell lung cancer (4), alteration of the metastatic potential of a human lung adenocarcinoma cell line (5) and reduction of metastasis in melanoma cells (6, 7). However, there is a paucity of work showing the effects of microgravity on the response of cancer cells to chemotherapeutic drugs. Hence, we posited the question whether simulated microgravity might alter the chemo-responsiveness of cancer cells in ways that affect cancer metastasis.

This question is important because metastasis leads to death in over 90% of cancer cases (8, 9). Metastasis is a complex multi-step process by which cancer cells spread from a primary cite to other locations in the body where they form secondary tumors. Metastasis occurs in all cancers, in spite of the very wide variety of cancers with respect to their molecular biology, pathogenesis and prognosis (10). Furthermore, the question is important because although chemotherapeutic drugs target and kill tumor cells during cancer treatment, emerging evidence suggests that some cancer drugs inadvertently promote metastasis (11–13). We recently showed that leukemic cancer cells treated with doxorubicin, an anti-cancer drug, migrated better than untreated cells, prior to cell death (14). Obviously, the ongoing search for anti-metastasis therapy (15) would benefit from a physical system that might alter any prometastatic effects of anti-cancer drugs, to improve therapeutic outcomes. We therefore hypothesized that microgravity might change the effects of cancer drugs on cellular functions involved in metastasis, such as migration.

To address our hypothesis, we considered the fact that disseminating tumor cells (DTCs) must be good at intravasating into and/or extravasating from blood vessels and migrating away from primary tumor cites in order to form new tumors (16, 17). Thus, we subjected cancer cells to 48 hours of microgravity and used standard migration assays to compare the migratory abilities of chemotherapy treated and untreated cancer cells, in order to assess whether microgravity alters our hitherto reported (14) inadvertent pro-metastatic effect of the anti-cancer drugs: daunorubicin (Dauno) and doxorubicin (Dox). Both Dox and Dauno are commonly used in the clinic against several cancers such as breast, lung and ovarian cancers, malignant melanomas and leukemia (18, 19). Interestingly, we find that post-microgravity chemotherapy using Dauno leads to increased chemotactic migration (p < 0.01) compared to normal gravity (1g) on earth in which there was reduced migration with Dauno. However, treatment of cells with doxorubicin led to enhanced migration both in 1g and following μg. Hence, microgravity modulates cancer cell response to chemotherapy in a drug-dependent manner, with possible impact for both space and terrestrial medicine.

## 2. Materials and Methods

### 2.1 Cell culture

We purchased both HL60 and K562 cell lines from ATCC (ATCC® CCL-240™ strain of HL60 and ATCC® CCL-243™ strain of K562). HL60 cells are multipotent promyelocytic leukemia cells derived from an acute myeloid leukemia (AML) patient (20, 21) which we have used extensively in previous studies involving cell differentiation, migration, cancer diagnostics, chemotherapy and radiotherapy (14, 22–25). The K562 cells are hematopoietic cancer cells, derived from a chronic myelogenous leukemia (CML) patient, that can differentiate into progenitors of erythrocytes, granulocytes and monocytes (26) (1). Both cell lines were cultured using RPMI 1640 (11875093, Life Technologies), supplemented with 10% (v/v) fetal bovine serum (FBS) and 1% Penicillin/Streptomycin as growth medium. They were grown in an incubator kept at 95% air; 5% CO_2_ and a temperature of 37°C. Cultures were maintained by the addition of fresh medium, between 1 × 10^5^ and 1.5 × 10^6^ cells/mL. Corning® T-75 flasks and T-25 flasks were used for sub-culturing. This required diluting the culture every 2 to 3 days. All experiments were carried out when cells were in the logarithmic growth phase. We chose these cell lines because current *in vivo* models of AML and CML based on the use of patient samples are difficult to establish and manipulate in the laboratory and both HL60 and K562 are still being used as *in vitro* models for assessing the efficacy of chemotherapeutic agents (27). Furthermore, HL60 and K562 have been used in microgravity studies (1, 28). Thus, our use of these cells enables the contextualization of our work for both terrestrial and space medical applications.

### 2.2 Simulation of microgravity

Simulated microgravity conditions were produced using a Rotary Cell Culture System (RCCS™) (Fig. 1) developed at the Johnson Space Center by NASA and made commercially available by Synthecon® Inc (Houston, TX, USA). The RCCS is a bioreactor equipped with 10-ml or 50-ml disposable High Aspect Ratio Vessel (HARV), with a silicon membrane on one side that provides gas exchange and designed to provide a low-shear cell culture system. Through vertical rotation, the RCCS produces solid-body rotation of the entire HARV and the cell culture medium, randomizing the gravity vector to simulate microgravity (29, 30). We seeded HL60 and K562 cells at a concentration of 2 × 10^5^ cells/ml in T-25 flasks (as static controls at normal gravity, 1g) and in 10-ml HARVs and rotated at 15 rpm (reported optimal rotation speed for many cells types including HL60 (28, 31) following the manufacturer’s protocols. Cells were kept in this microgravity condition for 48 hours, within a cell culture incubator (Fig. 1).

**Fig. 1.**
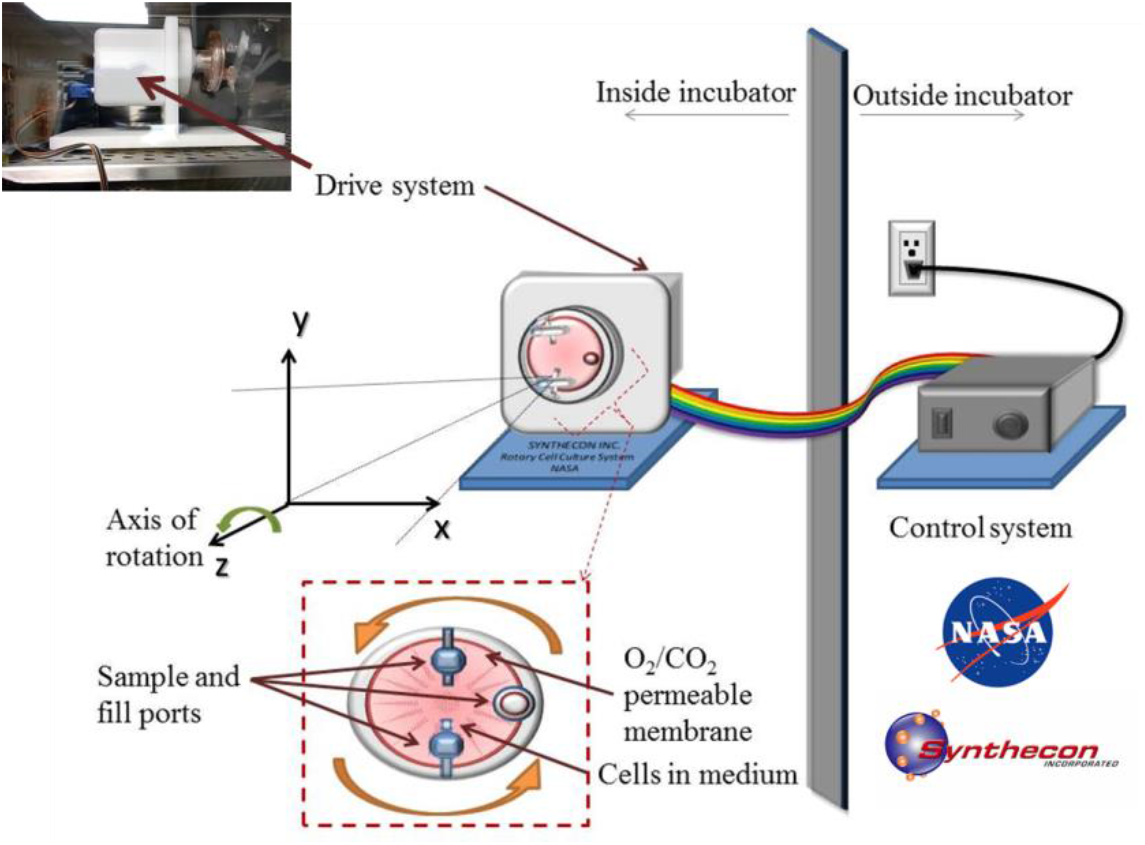
Simulation of microgravity inside a cell culture incubator using the rotary cell culture system, RCCS. The picture of the drive system as well as the schematic show the 10-ml cell culture vessel that is rotated to produce a time-averaged microgravity condition.

### 2.3 Chemotherapy and other drug treatments

Anthracyclines are among the main drugs used clinically against various cancers (13) and for leukemia, they are first line induction therapies (19, 32). The major anthracyclines are doxorubicin (Dox) (also known as adriamycin) and daunorubicin (Dauno) (also known as daunomycin). To closely replicate clinical dosage, we used Dox (Sigma 4458) at a final concentration of 5 μM (33, 34) and Dauno (Sigma D8809) at a final concentration of 1 μM (19, 32). To ascertain the role of F-actin and thereby attain molecular level insights, we used 2 μM Cytochalasin D (CytoD) to depolymerize F-actin and assess its well-established impact on migration (23, 24, 35).

### 2.4 Assessment of reactive oxygen species

To have molecular level insights into any changes caused by microgravity on cancer cells before chemotherapy, we assessed reactive oxygen species (ROS), using quantum dot fluorescence spectroscopy (25). ROS-sensitive signaling pathways are elevated in many kinds of cancer (36) and in prolonged microgravity (37).

### 2.5 Viability tests, morphometry and migration assay

We used Trypan blue exclusion to determine cell viability just before subjecting cells to microgravity and immediately after the 48 hours in microgravity. Morphometric data (size and shape) were extracted from phase contrast microscopy images of cells, using ImageJ, as described in our previous work (14). To quantify migration, we used the standard transwell cell migration and invasion assay (38) (Boyden chamber) with 5 μm pore size to ensure that cells actively squeeze through the polycarbonate membranes, as in 3D *in vivo* tissue, thereby mimicking the migratory phases of metastasis, including extravasation and intravasation. Following 48 hours of simulated μg, cells were treated with Dox and Dauno, placed in the transwell inserts and allowed to migrate chemotactically for 6 hours after which cells in the bottom plate and under the inserts were detached and counted. Care was taken to ensure that all conditions had the same cell density before the migration assay.

### 2.6 Measurement of cell deformability

In addition to migration, metastasis involves a circulatory phase in which cancer cells from primary tumor cites intravasate into the vasculature and are circulated around the body. In this circulatory metastatic phase, the short time-scale mechanical properties (24, 39) of such cells are important especially in capillaries (microvasculature) with internal diameters smaller than the size of cells at certain locations (40). We used our custom-developed microfluidic microcirculation mimetic MMM (14, 23, 40) to record the transit time of cells through 187 capillary-like constrictions (5 μm minimum diameter) and this transit time served as a readout of cell deformability.

### 2.7 Statistical analysis

All experiments were done at least three times (N1, N2 and N3). Averages were obtained for each experiment. Multiple experimental conditions (> 2) were compared for statistically significant differences using analysis of variance (ANOVA). Student T-Tests were used to compare two conditions. Analyses were done in OriginLab software (Northampton, Massachusetts, USA.)

## 3. Results

### 3.1 Cell viability and morphometry post-microgravity and post-microgravity chemotherapy

Following 48 hours of simulated microgravity (μg), cell viability remained between 98% and 100% for both static controls in normal gravity (1g) as well as for μg (Fig. 2A, 0 hr). The post-μg cells were immediately split into four flasks and treated with Dox, Dauno and Dox + CytoD and their viability was assessed and monitored every 2 hours (Fig. 2A). Since apoptosis does not set in until about 12 hours after chemotherapy and inadvertent prometastatic effects may only occur before this period (14) we monitored the viability every 2 hours until 6 hours post-μg (Fig 2A) and cell viability remained high as expected. In fact, at 6 hours, the reduction in cell viability was only statistically significant for the Dox+CytoD treated cells. Furthermore, using phase contrast microscopy and standard image segmentation algorithms for single cell morphometry, we found that in the first 6 hours post-μg chemotherapy, there was no statistically significant change in cell size except for the 3^rd^ trial for Dox+CytoD cells as shown in Fig 2B (see Fig S1 where trial N2 shows no significant difference). As previously obtained with chemotherapy in 1g (14), there is also with post-μg chemotherapy, a positive correlation in time between reduction in cell size and cell viability, a correlation indicative of increasing apoptosis. We found a similar trend for K562 cells.

**Fig. 2.**
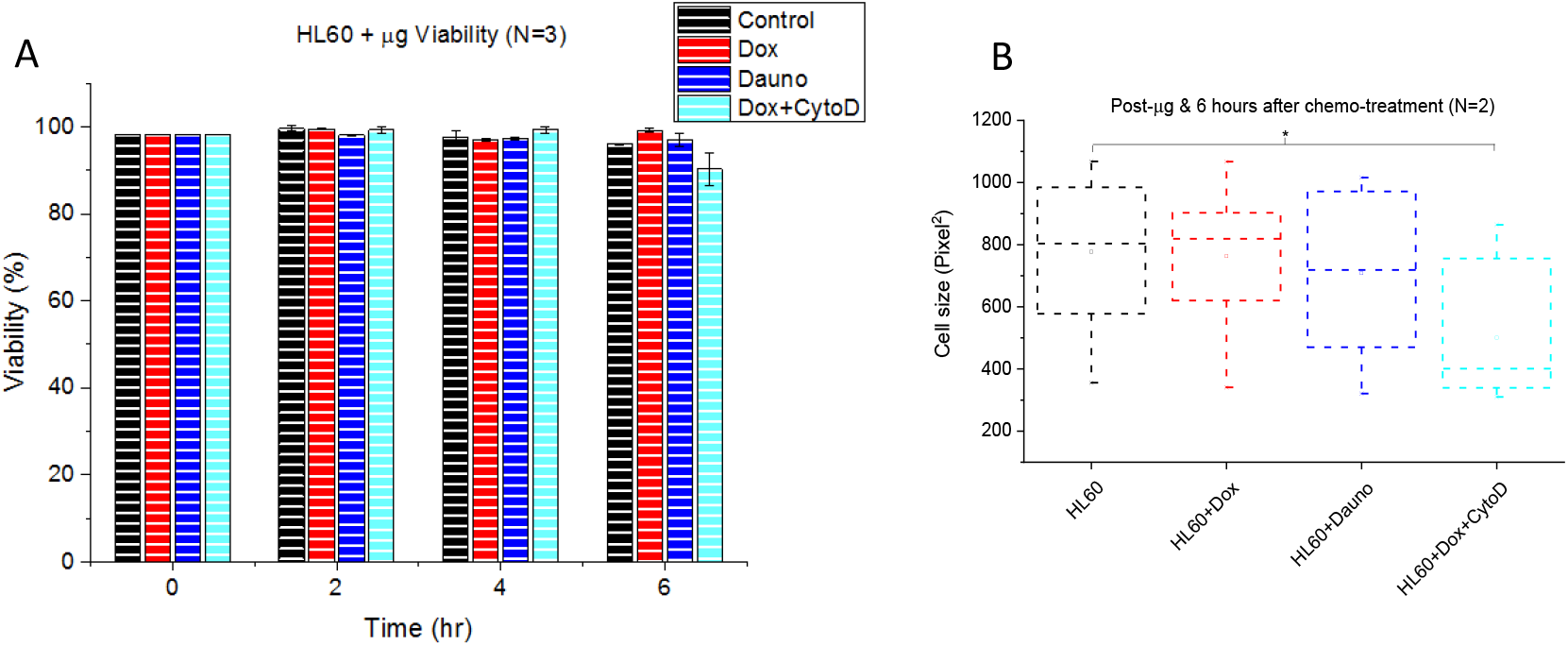
Cell viability/morphometry post-microgravity and post-microgravity chemotherapy. **(A)** Time-evolution of cell viability following microgravity (0 hr) and post-microgravity chemotherapy. At t = 6 hours, the viability was as follows: 95.9 ± 0.2% for HL60, 99.2 ± 0.3% for HL60+Dox, 96.9 ± 1.5% for HL60+Dauno, and 90.2 ± 3.8% for HL60+Dox+CytoD. The error bars are standard errors of the mean. (B). Box chart showing morphometric changes at 6 hours post-microgravity chemotherapy. Cells become significantly (p < 0.05) smaller in size after 6 hours of incubation with Dox (5 μM) and CytoD (2 μM).

### 3.2 Post-microgravity ROS generation is cell type dependent

Following our finding that 48 hours of simulated μg did not alter the viability of HL60 cells (Fig. 2A) and K562, we then assessed ROS in both HL60 and K562 cells to ascertain if known molecular level changes induced by μg also obtain in our setup, mainly as a further proof of the reliability of our setup and of the viability result (Fig. 2).

There was statistically no significant (NS) difference between the peak fluorescence intensity of QDs in a suspension of HL60 cells and QDs in a suspension of post-microgravity HL60 cells (Fig. 3A,B). This result was highly reproducible as can be seen in Fig. S2 showing N3, which is very comparable to N2 shown in Fig. 3A,B. However, for K562 cells, there was a statistically significant (p < 0.0001) reduction in peak fluorescence intensity between K562 cells in 1g and in μg (Fig. S3). Thus, microgravity-induced ROS generation is cell type dependent.

**Fig. 3.**
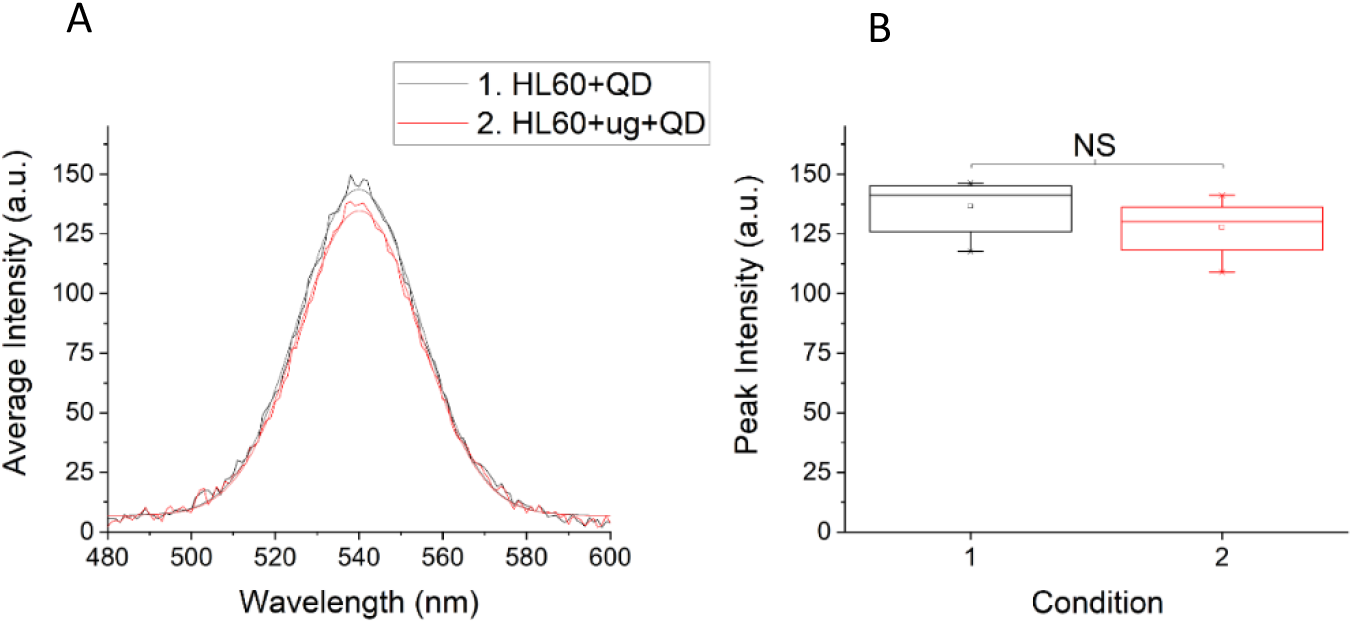
Assessment of ROS post-microgravity. (A) Quantum-dots fluorescence intensity peaks for HL60 cell suspension (HL60+QD) and post-microgravity HL60 cells (HL60+ug+QD). (B) Box chart comparing peak fluorescence intensities for the conditions in (A), showing non-significant (NS) difference based on T-Test.

### 3.3 Both doxorubicin and daunorubicin enhance post-microgravity migration of cells

Having found that 6 hours post-microgravity, cell viability remains above 95% even after treatment with chemotherapeutic drugs and that cell size does not change significantly (Fig. 2A,B) we focused on this 6-hour period since this is sufficient time for cells to extravasate from the vasculature and migrate into surrounding tissues, with metastatic consequences. Hence, we employed a chemoattractant based transwell migration assay to test how post-microgravity chemotherapy affects cell migration, a key step in the metastatic cascade. Just as we found for chemotherapy in 1g, doxorubicin significantly (p < 0.05) enhanced cell migration prior to inducing apoptosis, that is, 6 hours following induction (Fig. 4 A,B).

**Fig. 4.**
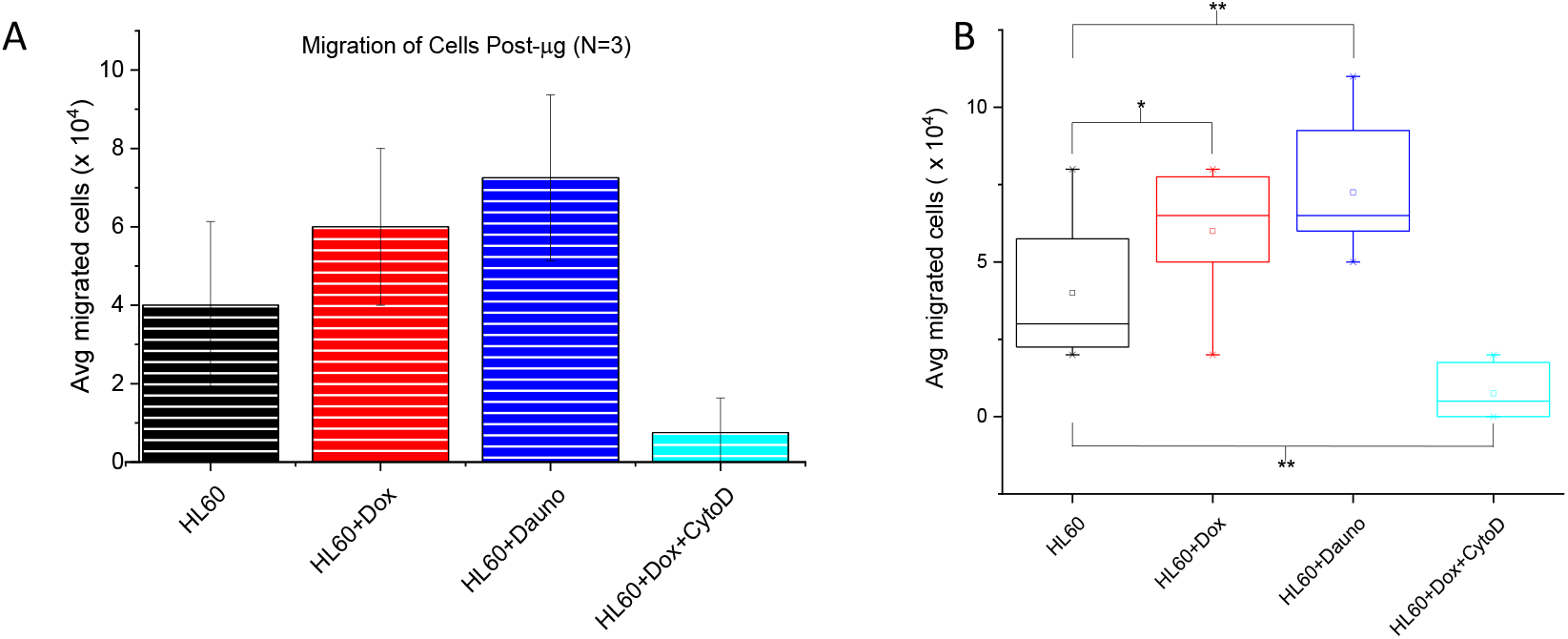
Post-microgravity chemotherapy alters rate of cancer cell migration due to F-actin reorganization. (A). Both 5 μM doxorubicin (Dox) and 1 μM daunorubicin (Dauno) enhance post-microgravity migration of cells. The reduced migration in CytoD treated cells reveals the dependence of the migration on F-actin organization. (B). Bar chart comparing the average migration rates of post-microgravity Dox and Dauno treated HL-60 cells with untreated. The chemotherapy-treated cells migrate significantly more (p < 0.05 for Dox and p < 0.01 for Dauno) than the untreated cells.

Since the determinant role of actin polymerization in cell migration is well established (23, 35), we depolymerized F-actin with 2 μM cytochalasin D (CytoD) in Dox treated cells and found as expected that these Dox+CytoD-treated cells did not migrate better than untreated HL60 cells. Surprisingly, post-microgravity chemotherapy using daunorubicin also led to a significant (p < 0.01) enhancement of the HL60 cell migration. This highly reproducible result (see Fig. S4 which is N2 while Fig. 4 is N3) is surprising because in 1g, we found and reported a significant reduction (p < 0.01) in migration of HL60 cells following daunorubicin treatment, compared to untreated cells. This finding ushers in the possibility of using microgravity as an immunomodulatory tool in the development of new cancer drugs with anti-metastatic capabilities.

With F-actin reorganization implicated in the chemotherapy-induced enhancement of migration, we investigated the effect of such reorganization on cell deformability using our microfluidic microcirculation mimetic (MMM) with transit times through the device as a surrogate for deformability. There was no clear correlation between transit times and enhanced migration (Fig. S5) leaving the connection between long time-scale migration (hours) and short time-scale changes in cell deformability (minutes and seconds) due to microgravity still an open question. Succinctly, we have found in this work a novel microgravity-induced modulation of chemotherapeutic effects on cell migration.

## 4. Discussion

Unlike plants which have special gravisensing cells called stomatocytes (41) the gravisensing cells in animals are yet to be discovered. Different animal cell types seemingly show different responses to microgravity. For instance, macrophages grown in simulated μg and later in the international space station (ISS) were recently found to respond to microgravity in seconds (42) confirming gravisensing by immune cells. Primary human macrophages grown in the ISS showed metabolic alterations and cytoskeletal stability (43). Both actin filaments and microtubules in endothelial cells were altered by simulated microgravity (44). Bone marrow stromal cells grown in simulated microgravity showed enhanced migration and neuroprotection following transplantation in a rat model for spinal cord injury (45). These recent findings resulted from over two decades of concerted efforts to grow cells and tissues in microgravity for use in space and terrestrial medicine (46, 47)

Here, we have shown that microgravity reverses the decreased cell migration obtained when cells were treated with daunorubicin in ground conditions (1g) (14), causing enhanced migration of HL60 cells following 48 hours of simulated μm. This intriguing result suggests interesting applications of simulated μm for both space and terrestrial medicine. For space medicine, our work posits that the therapeutic impact of certain drugs developed in 1g may not necessarily be replicated in μg environments and thus provides the rationale for space-based drug development beginning with *in vitro* μm conditions. For terrestrial medicine, the perennially urgent need for anti-metastasis drugs (15) and the recent success of immunotherapy (48) call for a synergy in which simulated μg is used as an immunomodulatory tool in the development of drugs that have both anticancer and anti-metastasis effects.

In our post-μg results, both daunorubicin and doxorubicin treatment lead to enhanced cell migration. Moreover, at comparable concentrations (doxorubicin, ⩾5 μM, and daunorubicin, ⩾1 μM) other researchers found in 1g conditions, increased generation of reactive oxygen species and a reorganization of the F-actin cytoskeleton (49) just as we have found in both 1g and μg. However, the reason(s) behind the marked differences between the two drugs with respect to 1g migration as well as the similar trend in μg are questions for further work. Molecular insights that might guide such further work would include the cell type dependence of the effect of microgravity on ROS generation which we have shown in this work. Along these lines, simulated microgravity was shown to increase ROS generation in mouse embryonic cells (37). Interestingly, the increased ROS generation in K562 cells which we have shown is consistent with the reported inhibition of K562 proliferation by simulated μg (4). With microgravity producing differences in ROS generation between a model of acute myeloid leukemia (AML), that is, HL60, and a model of chronic myelogenous leukemia (CML), that is, K562, which we have shown in this work, our suggested use of microgravity as an immunomodulatory tool for drug development becomes even more compelling.

## 5. Conclusions

Beginning with the hypothesis that microgravity might alter the effects of cancer drugs on cellular functions involved in metastasis, such as migration, we have shown in this work that indeed, post-microgravity chemotherapy can lead to results that are different from those of chemotherapy administered in normal gravity (1g). Daunorubicin treatment post-microgravity leads to enhanced cell migration within 6 hours of treatment, which is prior to cell death. Thus, microgravity reverses the effect of daunorubicin on cell migration prior to cell death. To the best of our knowledge, this is the first demonstration of the use of microgravity to modulate the effects of chemotherapeutic drugs on cancer cell migration. Furthermore, we found that microgravity modulates ROS generation in a cell-type dependent manner. These drug-specific and cell-specific modulations present microgravity as a cheap, non-invasive and easily accessible immunomodulatory tool for the development of anti-metastatic drugs against cancer as well as medication to enhance human space exploration.

## Acknowledgements

The authors are grateful to Dr Andrew Baruth and Dr Michael Nichols for allowing us the use of some of their laboratory equipment/supplies.

