## Supplementary Information for "Microgravity modulates effects of chemotherapeutic drugs on cancer cell migration"

**Cell viability and morphometry post-microgravity and post-microgravity chemotherapy**

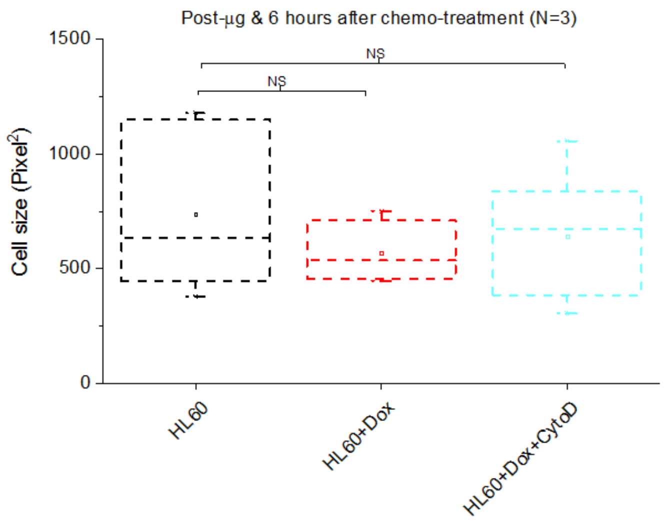


**Fig. S1. Box chart showing morphometric changes at 6 hours post-microgravity chemotherapy for trial 3 (N3).** Although cells become smaller in size after 6 hours of incubation with Dox (5 µM) and CytoD (2 µM) as in N2 (Fig. 2B) and N1, here, the reduction in size is not statistically significant (NS).

### Post-microgravity ROS generation is cell type dependent

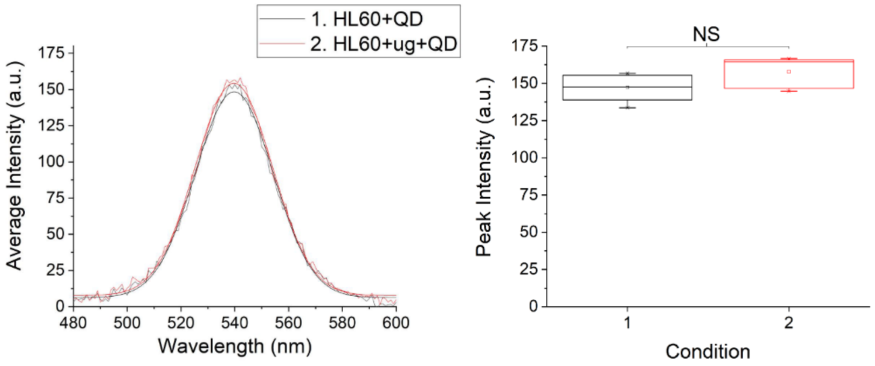


**Fig S2. Assessment of ROS post-microgravity.** (A) Quantum-dots fluorescence intensity peaks for HL60 cell suspension (HL60+QD) and post-microgravity HL60 cells (HL60+ug+QD). (B) Box chart comparing peak fluorescence intensities for the conditions in (A), showing non-significant (NS) difference based on T-Test.

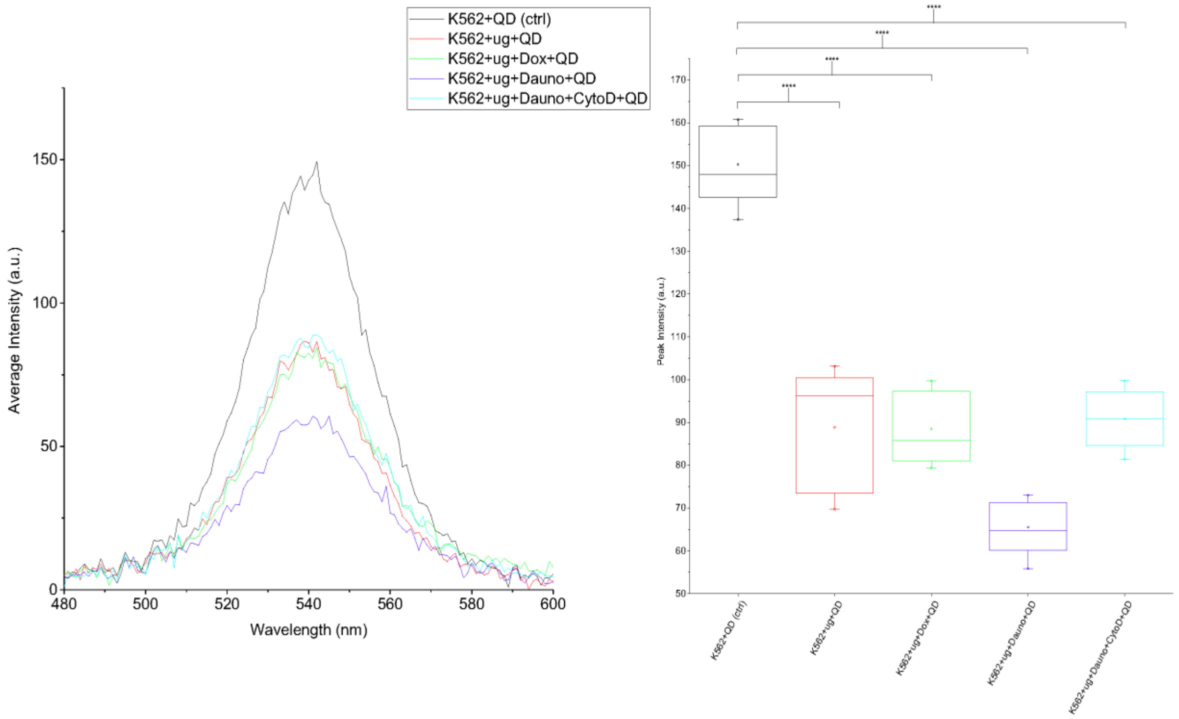


**Fig S3. Quantum dots fluorescence spectra and statistical comparison for K562 cells.** There are statistically significant differences ( $p < 0.0001$ for ****) between K562 cells in 1g and K562 cells following 48 hours of microgravity (K562+ug+QD), post-microgravity doxorubicin treatment (K562+ug+Dox+QD), daunorubicin treatment (K562+ug+Dauno+QD) and CytochalasinD treatment (K562+ug+Dox+CytoD+QD).

### Both doxorubicin and daunorubicin enhance post-microgravity migration of cells.

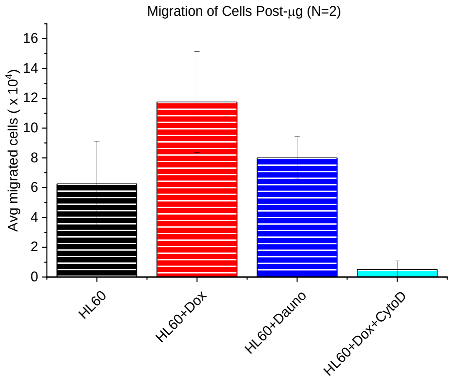


**Fig. S4 Post-microgravity chemotherapy alters rate of cancer cell migration due to F-actin reorganization, N2.** Both 5 μM doxorubicin (Dox) and 1 μM daunorubicin (Dauno) enhance post-microgravity migration of cells. The reduced migration in CytoD treated cells reveals the dependence of the migration on F-actin organization.

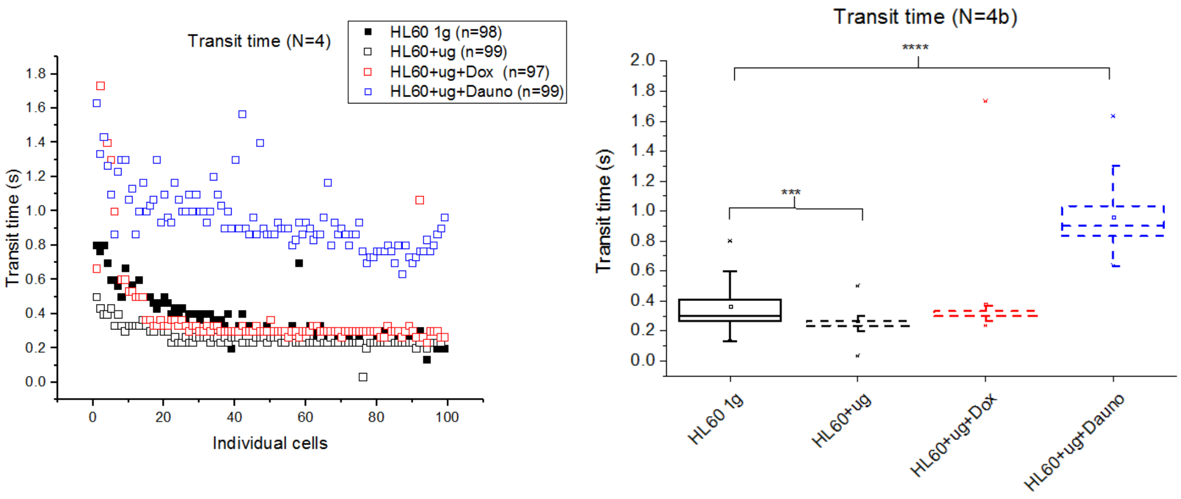


**Fig. S5. Transit times of cells advected through MMM.** (A) Scatter plots of transit times of individual cells comparing transit times of HL60 cells, post-microgravity HL60 cells (HL60+ug) and post-microgravity HL60 cells treated with doxorubicin (HL60+ug+Dox) or daunorubicin (HL60+ug+Dauno), 2 to 4 hours after treatment at a flow rate of (99µl/hr). (B) Box chart comparing transit times of cells described in (A). These results fluctuated and were not consistent between various trials (N1, N2, N3, N4 and N5). However, the various controls (HL60 in 1g) were consistent.
